# EmbB and EmbC Regulate the Sensitivity of *Mycobacterium abscessus* to Echinomycin

**DOI:** 10.1101/2024.02.25.578291

**Authors:** Jing He, Yamin Gao, Jingyun Wang, H.M. Adnan Hameed, Shuai Wang, Cuiting Fang, Xirong Tian, Jingran Zhang, Xingli Han, Yanan Ju, Yaoju Tan, Junying Ma, Jianhua Ju, Jinxing Hu, Jianxiong Liu, Tianyu Zhang

## Abstract

Treatment of *Mycobacterium abscessus* (Mab) infection is a major challenge due to its intrinsic resistance to most available drugs. It is thus imperative to find new anti-Mab drugs. In this study, we investigated the activity and intrinsic resistance mechanism of echinomycin (ECH) against Mab. ECH is active against Mab (MIC: 2 µg/mL). The *embC* gene knockout strain (Mab^ΔembC^) showed hyper-sensitive to ECH (MIC: 0.0078-0.0156 µg/mL). The MICs of ECH-resistant strains screened based on the Mab^ΔembC^ strain were 0.25-1 µg/mL. Mutations were found in the EmbB, including Asp306Ala, Asp306Asn, Arg350Gly, Val555Ile, and Gly581Ser, which led to increased resistance to ECH when overexpressed in Mab^ΔembC^ individually (0.25-0.5 µg/mL). The EmbB mutants edited by the CRISPR/Cpf1 system became more resistant to ECH (MIC: 0.25-0.5 µg/mL). The permeability of gene-edited and overexpressed Mab strains was reduced, as shown by the ethidium bromide accumulation assay, but it was still significantly higher than that of the parent Mab. To summarize, our study demonstrates that ECH has a strong anti-Mab activity and confirms that EmbB and EmbC are related to the sensitivity of Mab to ECH. EmbB mutation may partially compensate for the function of EmbC.

**Impact Statement:** *Mycobacterium abscessus* (Mab) is a rapidly growing, intrinsic multidrug-resistant Mycobacterium. This study demonstrated that echinomycin (ECH) has potent antibacterial activity against Mab, and the mechanism of ECH resistance to Mab is related to EmbB and EmbC. EmbB and EmbC can alter the sensitivity of Mab to ECH by altering the permeability of its cell wall. In addition, there is a functional complementary evolution between EmbB and EmbC to regulate sensitivity to ECH. Overall, our study provides a novel anti-Mab drug candidate ECH and a scientific foundation for developing effective strategies to prevent and control Mab.

## INTRODUCTION

Infections caused by non-tuberculous mycobacteria (NTM) are increasing worldwide^1^. The diseases associated with NTM are extremely challenging to treat because these bacteria are naturally resistant to many conventional antibiotics^2^. NTM are diverse and ubiquitous in the environment, with only a few species causing serious and often opportunistic infections in humans, including *Mycobacterium abscessus* (Mab)^3, 4^. This rapidly growing mycobacterium is one of the most commonly identified NTM species causing severe respiratory, skin, and mucosal infections in humans^5^. It is often regarded as one of the most antibiotic-resistant mycobacteria, leaving us with few therapeutic options^6^. Therefore, a detailed understanding of resistance mechanisms and developing potent anti-Mab drugs for therapeutic use is urgently needed.

The cell wall serves as the interface between the external environment and the internal cellular components, holding a responsibility to perform many important biological functions, including structural integrity, protection, and transport. Cell walls are crucial for the survival of bacteria, therefore, many enzymes involved in bacterial cell wall biosynthesis are gaining attention as potential targets for developing novel drugs. The mycobacterial *embCAB* operon encodes arabinosyltransferases, which are involved in the biosynthesis of arabinogalactan (AG) and lipoarabinomannan (LAM)^7, 8^. AG and LAM are key components of the mycobacterial cell wall^9, 10^. The EmbA and EmbB proteins are involved in synthesizing the galactan and mannan regions of AG and LAM^11^. They recognise and transfer galactose and mannose sugar molecules onto the growing polysaccharide chains^12^. The EmbC is a transmembrane protein that plays a crucial role in extending the arabinan chains in AG and LAM^13^. Ethambutol (EMB), a first-line anti-tuberculosis drug, can mediate the addition of arabinose units to the growing polysaccharide chains, thereby determining the length and structure of the arabinan^14^. Inhibition of arabinan synthesis eliminates the mycolic acid anchoring site, causing a reduction in the accumulation of mycolic acids in the cell wall. Ultimately, this damages the cell wall’s integrity and leads to cell death^15, 16^. Mutations in the *embC*, *embB*, and *embA* genes have been associated with *Mycobacterium tuberculosis* (Mtb) resistance to the EMB^17^. The mechanism of Mab’s natural resistance to EMB is related to polymorphisms in the *embB* gene, specifically the substitution of isoleucine by glutamine at position 303 and leucine by methionine at position 304 of the arabinosyltransferase EmbB^18^. Recently, our lab reported that knockout of the *embC* gene can synthesize only LM but not LAM, leading to increased cell membrane permeability and making Mab susceptible to many originally ineffective drugs^19^, the specific mechanism behind this is not yet clearly explained in Mab. Studies have shown a similar explanation in *Mycobacterium smegmatis* that the absence of LM is not crucial for maintaining capsule integrity, while the lack of LAM can weaken the structural integrity of peptidoglycan cell walls, causing them to lose structural stability and begin to develop “bubble-like beads”, indicating that the cell wall is no longer firm and unable to maintain cell shape to resist against swelling pressure^20^. Taken together, these suggest that EmbCAB proteins are good targets for developing antimycobacterial drugs.

Recently, it was reported that echinomycin (ECH) has antimycobacterial effect against Mtb H37Ra (Minimum inhibitory concentration, MIC: 0.5 µg/mL) and *Mycobacterium bovis* (M. bovis) (MIC: 0.1 µg/mL)^21^. However, the mechanism of action and activity of ECH against NTM has yet to be fully explored. In this study, we found that ECH has potent antibacterial activity against Mab (MIC: 2 µg/mL), and its resistance mechanism is related to the *embCAB* operon. Our findings certificated that EmbB and EmbC are related to the sensitivity of Mab to ECH and provided a further hypothesis that the functional complementary evolution of EmbB and EmbC leads to changes in the sensitivity of Mab to ECH.

## RESULTS

### Sensitivities of AlMab and Mab**^Δ^**^embC^ to ECH

The MIC_lux_ of ECH against autoluminescent Mab (AlMab) was determined by detecting RLUs (Relative light units). MIC_lux_ of ECH against AlMab was defined as the lowest concentration inhibiting more than 90% RLUs compared to that of the drug-free control. ECH showed an inhibitory effect on AlMab in a concentration-dependent manner with a MIC of 2 µg/mL (Figure 1). The sensitivity of ECH against the *embC-*ko (gene knock out) Mab strain (Mab^ΔembC^) was determined by the broth dilution method. MIC_liquid_ is the lowest concentration observed with the naked eye without bacterial growth. Observing the transparent 96-well plate, the *in vitro* MIC_liquid_ of Mab^ΔembC^ treated with ECH ranged from 0.0078 to 0.0156 μg/mL.

**Figure 1.**
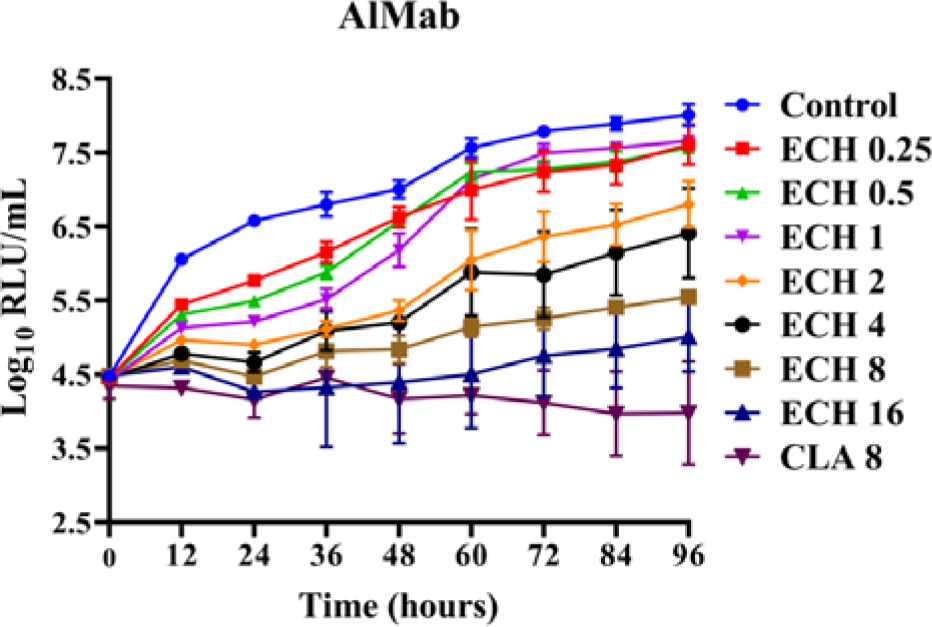
Time-killing curves of ECH against AlMab in liquid culture. Control: DMSO, Dimethyl sulfoxide; ECH: Echinomycin; CLA: Clarithromycin. Numbers after the drugs indicating the concentrations tested (μg/mL); Data are expressed as mean ± SD from three independent biological repeats. The experiment was performed in triplicate (three independent experiments), and the representative results are shown.

### Mutation sites of Mab**^Δ^**^embC^ mutant strains resistant to ECH

Among the screened Mab^ΔembC^ spontaneous drug-resistant mutant strains, 6 strainswere randomly selected to perform whole-genome sequencing (WGS) with their parent strain (Mab^ΔembC^) as a control. The WGS results showed that the *embB* was the main mutated gene in these mutant strains. To confirm the mutations, the *embB* was amplified by PCR and sequenced at Sangon Biotech (Shanghai). We identified four mutations in *embB* gene of Mab^ΔembC^ as shown in Table 1.

**Table 1.**
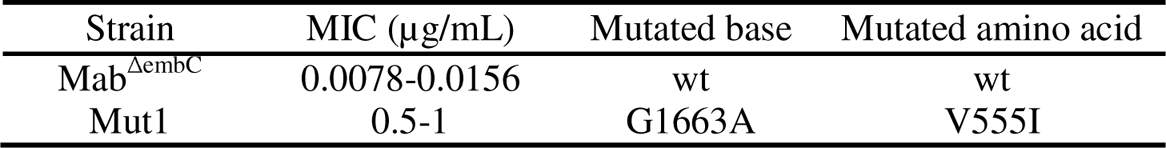
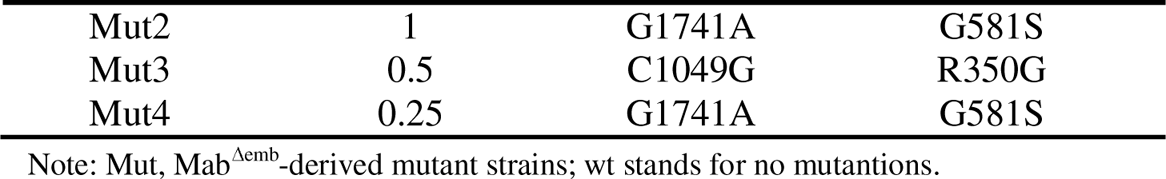
Sensitivity of ECH against the Mab^ΔembC^ mutant strains and mutations in *embB* by WGS.

| Strain | MIC ($\mu\text{g/mL}$) | Mutated base | Mutated amino acid |
| --- | --- | --- | --- |
| Mab<sup>$\Delta\text{embC}$</sup> | 0.0078-0.0156 | wt | wt |
| Mut1 | 0.5-1 | G1663A | V555I |
| Mut2 | 1 | G1741A | G581S |
| Mut3 | 0.5 | C1049G | R350G |
| Mut4 | 0.25 | G1741A | G581S |

Note: Mut, Mab<sup>Δemb</sup>-derived mutant strains; wt stands for no mutations.

The *embB* genes of the remaining mutants were amplified by PCR. Their PCR products were sequenced, and the results are shown in Table 2. Among the 11 ECH-resistant mutants, 7 strains had mutations in the *embB* gene different from those found by WGS; 4/7 of them had mutations at nucleotide position 916 of the *embB* gene, resulting in Asp306Asn amino acid substitution in EmbB, while 3/7 strains had mutations at nucleotide position 917 of the *embB* gene, resulting in Asp306Ala amino acid substitution in EmbB. However, 1 strain was detected without any mutation in *embB* and its upstream sequence (∼200 bp), which indicated that other ECH resistance mechanisms may exist.

**Table 2.** MICs of ECH to the Mab^ΔembC^ mutant strains and mutations in *embB* detected directly by PCR.

| Strain | MIC ( μg/mL ) | Mutated base | Mutated amino acid |
| --- | --- | --- | --- |
| Mab<sup>ΔembC</sup> | 0.0078-0.0156 | wt | wt |
| Mut7<sup>a</sup> | 0.5 | G916A | D306N |
| Mut8 | 1 | C1048G | R350G |
| Mut9<sup>a</sup> | 0.5 | G916A | D306N |
| Mut10<sup>a</sup> | 1 | G916A | D306N |
| Mut11<sup>a</sup> | 1 | G916A | D306N |
| Mut12 | 0.25 | wt | wt |
| Mut13 | 1 | C1048G | R350G |
| Mut14 | 0.5 | C1048G | R350G |
| Mut15<sup>b</sup> | 0.5 | A917C | D306A |
| Mut16<sup>b</sup> | 0.25 | A917C | D306A |
| Mut17<sup>b</sup> | 0.25 | A917C | D306A |

Note: <sup>a</sup> and <sup>b</sup> indicate new mutations different from those found in strains listed in Table 1 by WGS; wt stands for no mutations.

According to the above sequencing results, a total 5 different point mutations were found in the *embB* gene of Mab^ΔembC^ (Table 3)

**Table 3.** Mutation types of Mab-*embB* in Mab^ΔembC^.

| Number | Mutation types |  |
| --- | --- | --- |
|  | Base mutation | Amino acid mutation |
| Mut1 | G1663A | V555I |
| Mut2 | G1741A | G581S |
| Mut3 | A917C | D306A |
| Mut4 | C1049G | R350G |
| Mut5 | G916A | D306N |

Note: Mut stands for mutation types.

### MICs of different antibiotics against Mab**^Δ^**^embC^ mutant strains

The broth dilution method was used to detect the MICs of different antibiotics (moxifloxacin (MXF), levofloxacin (LEV), vancomycin (VAN), rifampin (RIF), rifabutin (RFB), ethambutol (EMB), linezolid (LZD), amikacin (AMK), cefoxitin (CEF) and clofazimine (CLF)) to the Mab^ΔembC^-derived ECH-resistant mutants (Table 4). The mutants became more resistant to ECH than Mab^ΔembC^, and the MICs of some antibiotics, such as VAN and RIF, against the Mab^ΔembC^ mutants also changed.

**Table 4.**
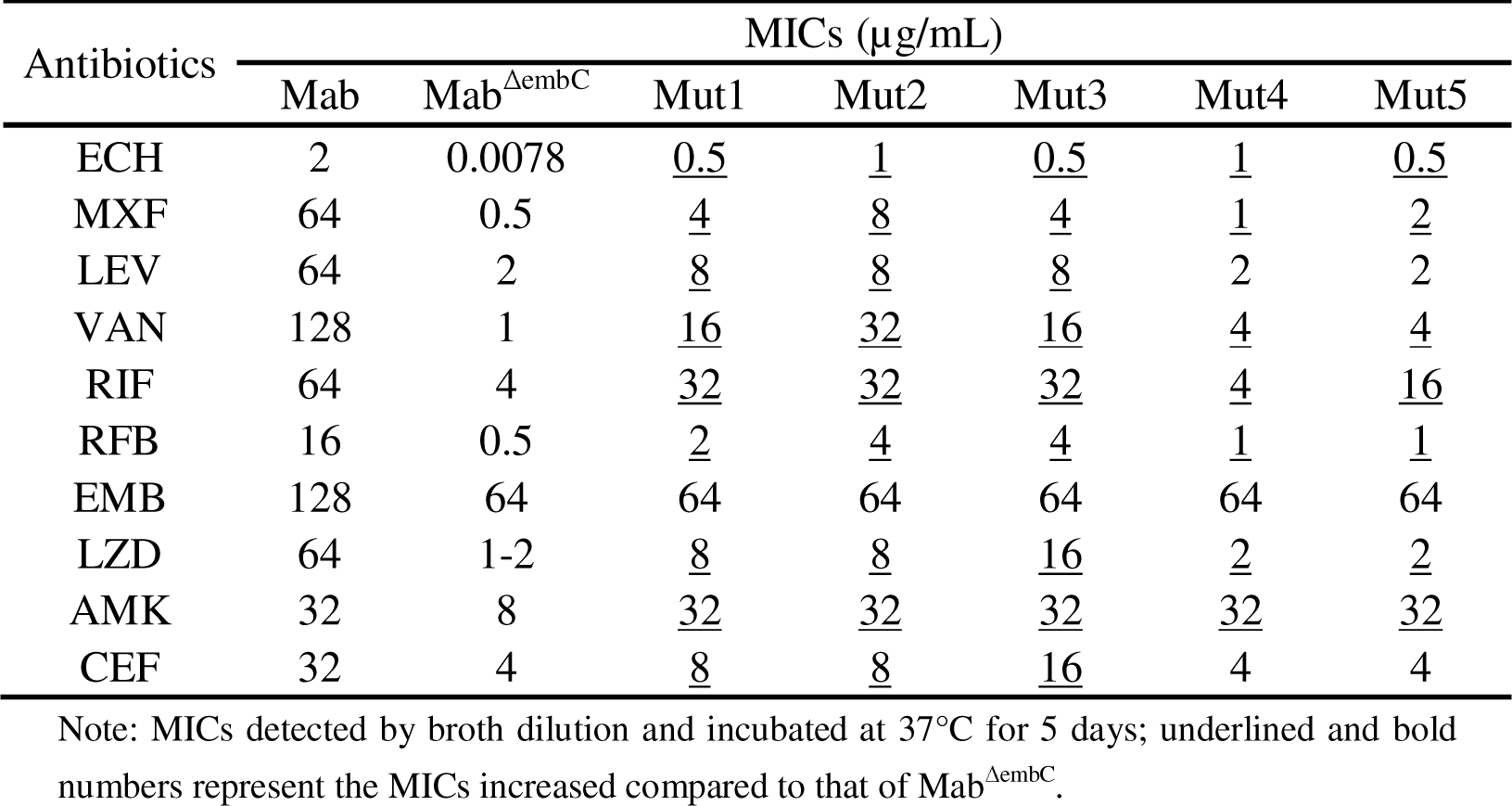
MICs of different antibiotics against Mab^ΔembC^ mutant strains.

### Sensitivities of Mab**^Δ^**^embC^ overexpressing wild-type and mutant *embB* genes to different antibiotics

We found that the strain overexpressing the wild-type *embB* gene has no significant change in drug sensitivity to various antibiotics, including ECH, while the other strains, which overexpress the mutant *embB* gene, become more resistant to ECH (Table 5). The changes in MICs of other antibiotics to all the overexpression mutant strains were ≤ 4-fold except for the MIC of RIF to Δpmut2 (*embB^G^*^581^*^S^* overexpressed in Mab^ΔembC^) strain.

**Table 5.** Drug sensitivities of overexpression strains to different antibiotics ntibiotics.

| Antibiotics | MICs ( μg/mL ) | | | | | | | |
| --- | --- | --- | --- | --- | --- | --- | --- | --- |
| | $\Delta$ | Δpmut1 | Δpmut2 | Δpmut3 | Δpmut4 | Δpmut5 | Δpwt | ΔpV |
| ECH | 0.0156 | 0.25 | 0.5 | 0.125 | 0.125 | 0.5 | 0.0625 | 0.0625 |
| LEV | 2 | 2 | 4 | 2 | 4 | 8 | 2 | 2 |
| VAN | 1 | 2 | 2 | 2 | 4 | 4 | 1 | 1 |
| RIF | 4 | 4 | 64 | 8 | 8 | 8 | 2 | 2 |
| RFB | 0.5 | 0.25 | 1 | 2 | 2 | 1 | 0.5 | 1 |
| EMB | 64 | 64 | 128 | 64 | 64 | 64 | 64 | 32 |
| LZD | 1 | 1 | 2 | 1 | 2 | 4 | 1 | 1 |
| AMK | 8 | 8 | 8 | 32 | 8 | 8 | 4-8 | 16 |
| CEF | 4 | 8 | 8 | 16-32 | 8 | 32 | 8 | 8 |
| MXF | 0.5 | 0.5 | 1 | 1 | 1 | 2 | 0.5 | 0.5 |

Note: $\Delta$ means Mab <sup>$\Delta$ embC</sup>, *embC* gene knockout Mab; $\Delta$ pmut1, *embB*<sup>V551I</sup> overexpressed in Mab <sup>$\Delta$ embC</sup>; $\Delta$ pmut 2, *embB*<sup>G581S</sup> overexpressed in Mab <sup>$\Delta$ embC</sup>; $\Delta$ pmut3, *embB*<sup>D306A</sup> overexpressed in Mab <sup>$\Delta$ embC</sup>; $\Delta$ pmut4, *embB*<sup>R350G</sup> overexpressed in Mab <sup>$\Delta$ embC</sup>; $\Delta$ pmut5, *embB*<sup>D306N</sup> overexpressed in Mab <sup>$\Delta$ embC</sup>; $\Delta$ pwt, *embB*<sup>wt</sup> overexpressed in Mab <sup>$\Delta$ embC</sup>; $\Delta$ pV, Mab <sup>$\Delta$ embC</sup> with pMV261A'-the empty vector. MICs detected by broth dilution after incubation at 37°C for 5 days.

### Drug sensitivities of *embB* gene-edited Mab^ΔembC^ strains

The drug susceptibility testing of successfully edited strains to ECH and other antibiotics showed that in addition to the significant rise in MIC to ECH, the MIC of VAN against edited Mab^ΔembC^ was also increased. VAN is a glycopeptide antibiotic that inhibits cell wall synthesis.

**Table 6.** MICs of ECH and other antibiotics to *embB*-edited strains Antibiotics.

| Antibiotics | MICs ( μg/mL ) | | | | | |
| --- | --- | --- | --- | --- | --- | --- |
| | $\Delta$ | EΔ-V555I | EΔ-G581S | EΔ-D306A | EΔ-R350G | EΔ-D306N |
| ECH | 0.0156 | 0.25 | 0.5 | 0.25 | 0.25 | 0.5 |
| MXF | 0.5 | 4 | 4 | 2 | 2 | 4 |
| LEV | 2 | 8 | 2 | 4 | 4 | 4 |
| <u>VAN</u> | 1 | 8 | 4 | 32 | 4 | 4 |
| RIF | 4 | 8 | 16 | 16 | 16 | 16 |
| RFB | 0.5 | 2 | 2 | 2 | 1 | 2 |
| EMB | 64 | 64 | 32 | 32 | 16 | 64 |
| LZD | 1-2 | 4 | 2 | 4 | 8 | 4 |
| AMK | 8 | 16 | 8 | 16 | 16 | 8 |
| CEF | 4 | 4 | 8 | 8 | 8 | 16 |

Note: $\Delta$ means $\text{Mab}^{\Delta\text{embC}}$ ; E means *embB*-edited $\text{Mab}^{\Delta\text{embC}}$ strains; MICs detected by broth dilution after incubation at 37°C for 5 days.

### Detection of the sensitivities of different strains to ECH on plates

The results comparing the sensitivities of the *embB* overexpressed Mab^ΔembC^ strains and Mab^ΔembC^ to ECH are shown in Figure 2. The growth status of various strains on the drug-free 7H10 plate is the same, but as the concentration of ECH increases, compared with Mab^ΔembC^, the *embB* overexpressed Mab^ΔembC^ showed more resistant to ECH.

**Figure 2.**
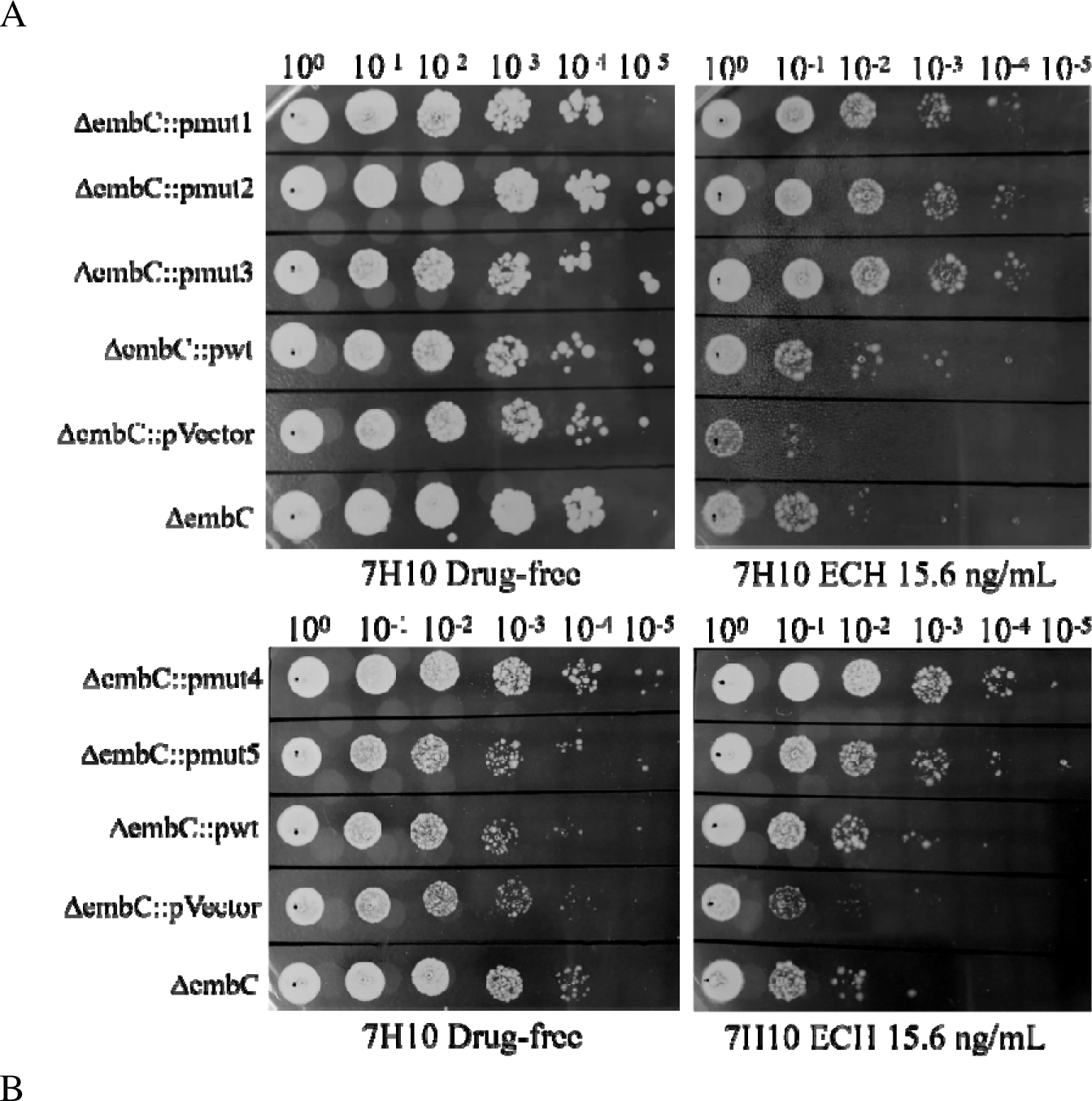
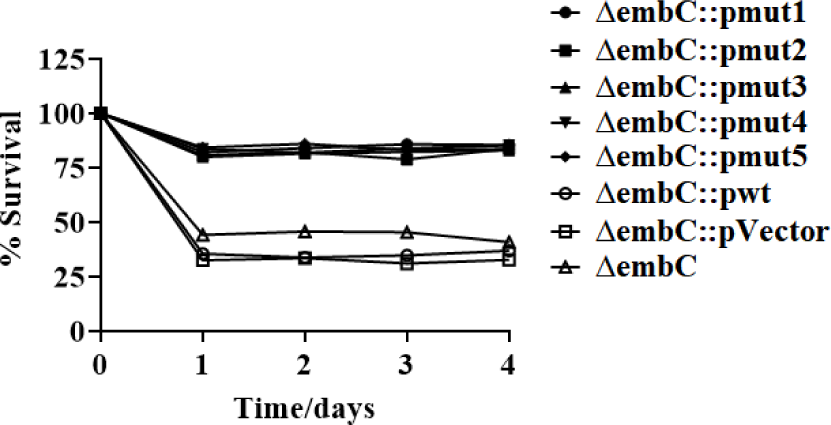
**A)** Sensitivities of the overexpressed strains to ECH on plates. ΔembC, Mab^ΔembC^, *embC* gene knockout Mab; ΔembC::pmut1, *embB*^V555I^ overexpressed in Mab^ΔembC^; ΔembC::pmut2, *embB^G^*^581^*^S^* overexpressed in Mab^ΔembC^; ΔembC::pmut3, *embB^D^*^306^*^A^* overexpressed in Mab^ΔembC^; ΔembC::pmut4, *embB^R^*^350^*^G^* overexpressed in Mab^ΔembC^; ΔembC::pmut5, *embB^D^*^306^*^N^* overexpressed in Mab^ΔembC^; ΔembC::pwt, *embB^wt^* overexpressed in Mab^ΔembC^; ΔembC::pVector, Mab^ΔembC^ with pMV261A’. These strains were serially diluted, spotted onto 7H10-agar plates with or without ECH and incubated at 37°C for 7 days. **B)** Bacterial survival rate after exposure to ECH (15.6 ng/mL). All data are representative of three independent experiments.

The results comparing the sensitivities of the *embB*-edited Mab^ΔembC^ strains and Mab^ΔembC^ to ECH are shown in Figure 3. Compared with Mab^ΔembC^, the edited Mab^ΔembC^ strains became more resistant to ECH on the plates, indicating that the mutation of *embB* indeed changed the sensitivity of Mab^ΔembC^ to ECH.

**Figure 3.**
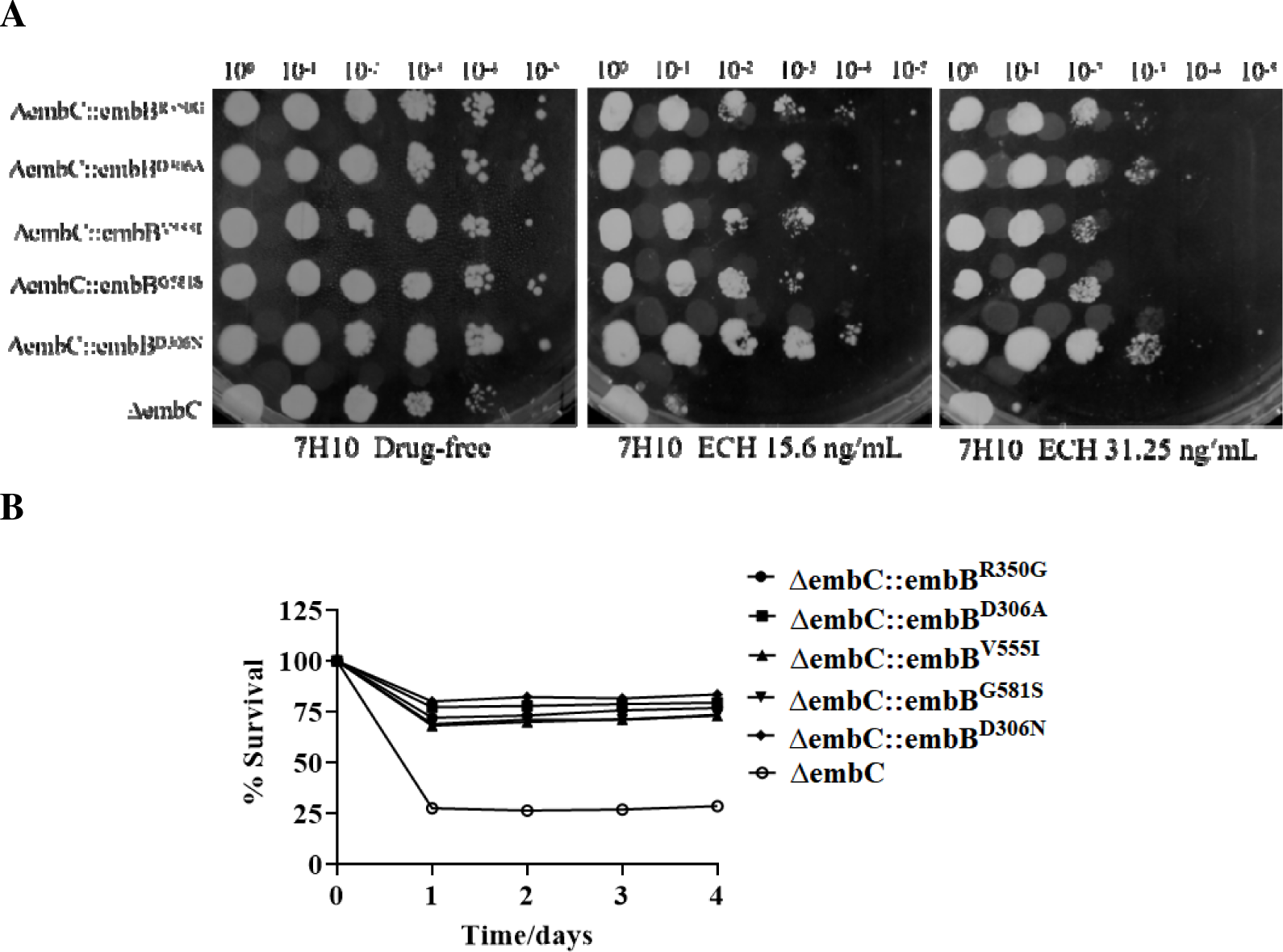
**A)** Sensitivity of edited strains to ECH on plates. ΔembC, Mab^ΔembC^, *embC* gene knockout Mab; ΔembC::embB^R350G^, *embB^R^*^350^*^G^* edited Mab^ΔembC^; ΔembC::embB^D306A^, *embB^D^*^306^*^A^* edited Mab^ΔembC^; ΔembC::embB^V555I^, *embB^V^*^555^*^I^* edited Mab^ΔembC^; ΔembC::embB^G581S^, *embB^G^*^581^*^S^* edited Mab^ΔembC^; ΔembC::embB^D306N^, *embB^D^*^306^*^N^* edited Mab^ΔembC^; These strains were serially diluted, spotted onto 7H10-agar plates with or without ECH and incubated at 37°C for 7 days. **B)** Bacterial survival rate after exposure to ECH (31.25 ng/mL). All data are representative of three independent experiments.

Comparing the sensitivities of strains with the same *embB* mutation but obtained from different routes to ECH on the plates: A *embB^R^*^350^*^G^* was selected to compare the sensitivities of the spontaneously resistant mutant and *embB-*edited strain, as well as the strains overexpressing wild-type or mutant *embB* in Mab^ΔembC^. As shown in Figure 4, the growth of various strains on 7H10 plates without drugs were the same. But as the concentration of ECH increases, all the Mab strains containing the *embB^R^*^350^*^G^* gene showed more resistant to ECH on the plates, indicating that the mutation of *embB* indeed changes the sensitivity of Mab^ΔembC^ to ECH.

**Figure 4.**
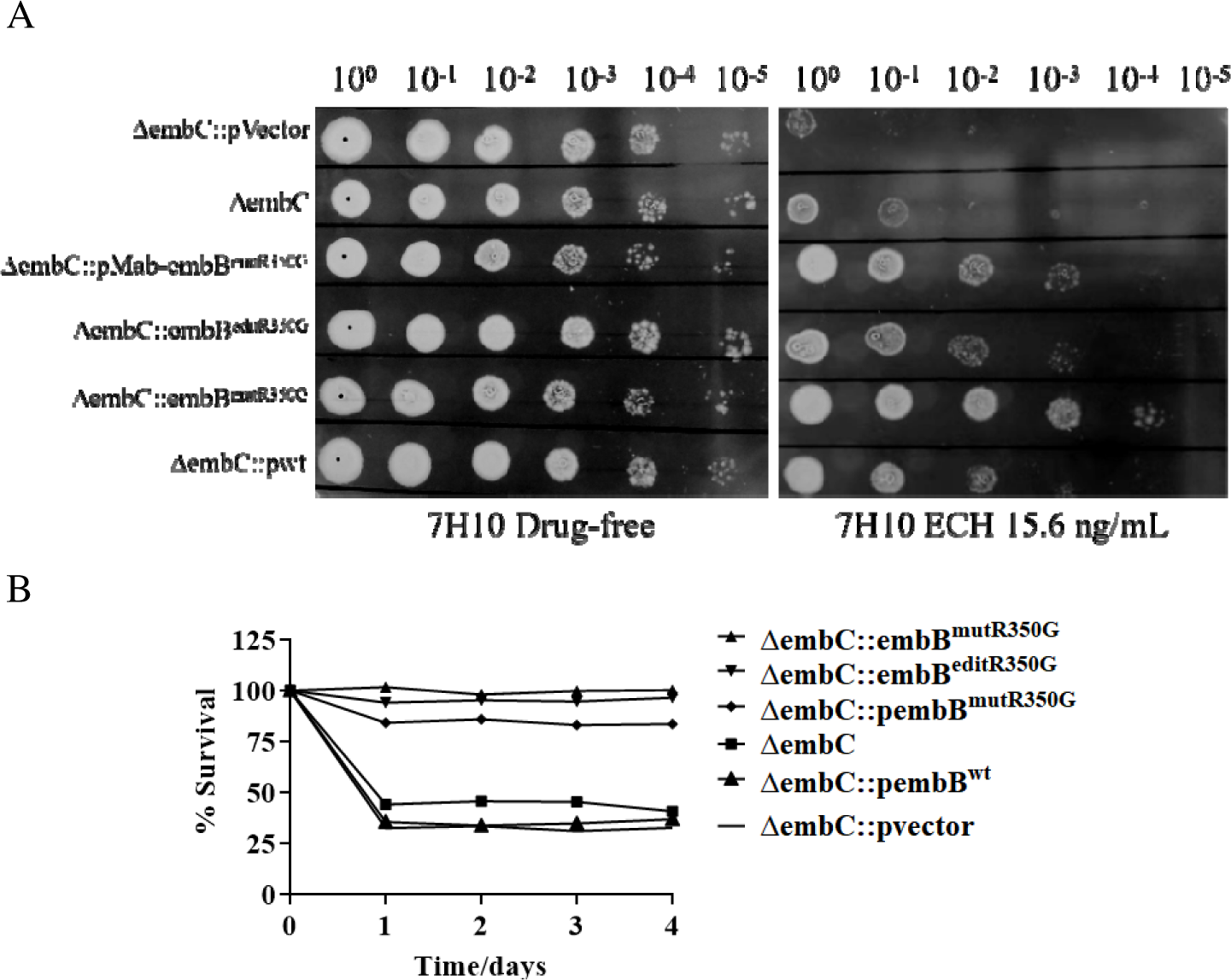
**A)** Comparing sensitivities of different Mab strains to ECH on plates. ΔembC, Mab^ΔembC^, *embC* gene knockout Mab; ΔembC::pVector, Mab^ΔembC^ with pMV261A’; ΔembC::pMab-embB^mutR350G^, *embB^R^*^350^*^G^* overexpressed in Mab^ΔembC^; ΔembC::embB^editR350G^, *embB^R^*^350^*^G^* edited Mab^ΔembC^; ΔembC::embB^mutR350G^, *embB^R^*^350^*^G^* spontaneous mutant Mab^ΔembC^; ΔembC::pwt, *embB^wt^*overexpressed in Mab^ΔembC^; These strains were serially diluted, spotted onto 7H10-agar plates with or without ECH and incubated at 37°C for 7 days. **B)** Bacterial survival rates after exposure to ECH (31.25 ng/mL). All data are representative of three independent experiments.

### Cell wall permeability results

Ethidium bromide (EtBr) can enter the cell through damaged cell walls and bind to nuclear DNA, emitting red-orange fluorescence under ultraviolet light. The fluorescence increases when combined with double-stranded DNA. The EtBr accumulation experiment uses EtBr as a substrate. By detecting its fluorescence value after entering the cell, the accumulation amount in the cell can be visualized to tell the damage to the cell wall.

The cell wall permeability results illustrated that compared to Mab^ΔembC^, the accumulation of EtBr in spontaneous ECH-resistant mutant Mab^ΔembC^ strains, mutated *embB* overexpressed Mab^ΔembC^ and edited Mab^ΔembC^ was obviously reduced. However, it was still higher than the accumulation of *embC* complemented Mab^ΔembC^ or Mab^wt^ (Figure 5). While the EtBr accumulation of overexpressing wild-type *embB* in Mab^ΔembC^ showed no significant change relative to Mab^ΔembC^. We further compared the sensitivity of these strains to crystal violet. As previous reports, crystal violet, peacock green, and SDS are commonly used to evaluate the toxicity of mycobacteria to determine the cell walls permeability^22^. The results are consistent with the accumulation experiment of ethidium bromide. The mutated EmbB could possibly compensate for the partially missing function of EmbC to reduce the permeability of the Mab^ΔembC^ cell wall. In other words, the mutated EmbB can compensate for the part of the EmbC function to restore part of the cell wall integrity. However, it cannot fully replace the EmbC.

**Figure 5.**
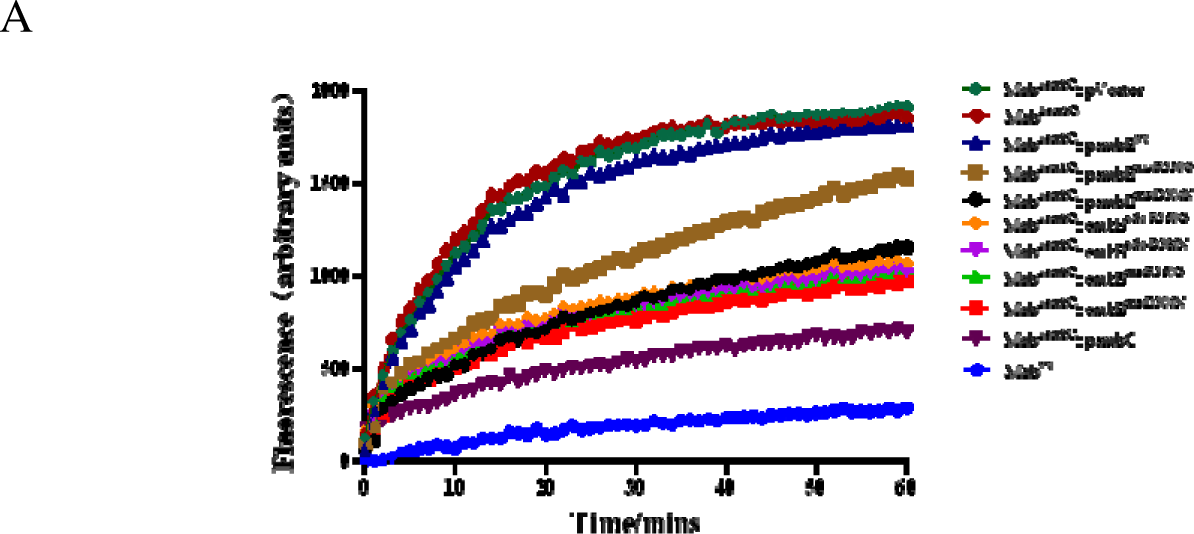
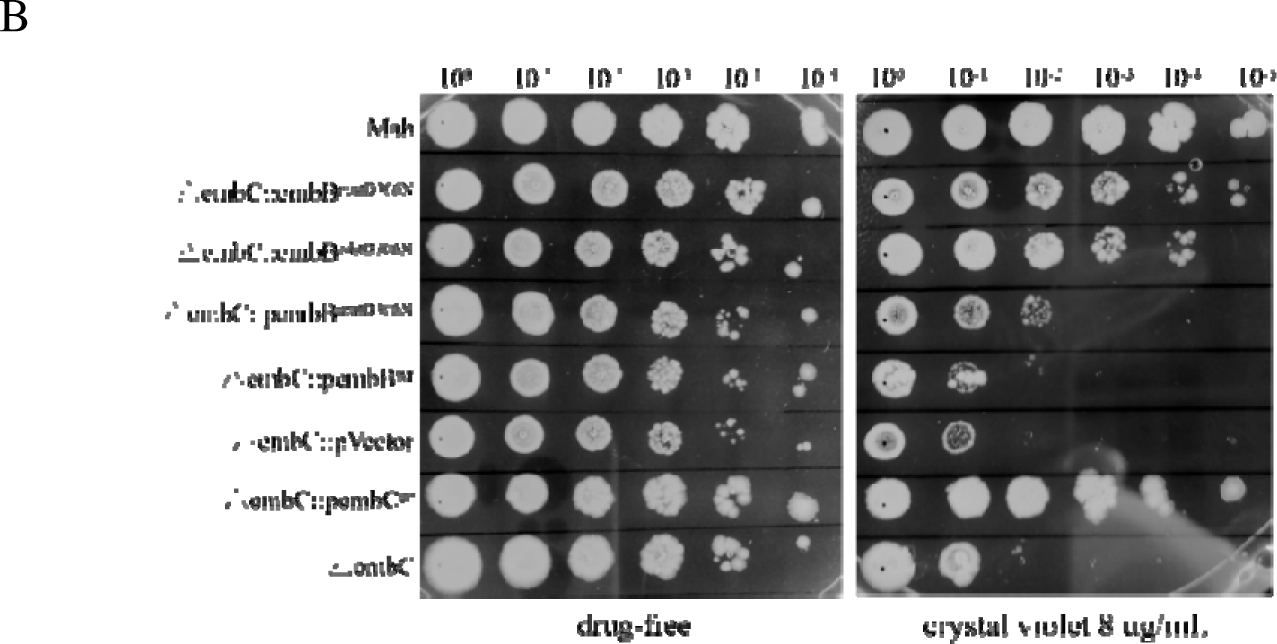
**A)** Accumulation of EtBr in different strains. Mab^ΔembC^, *embC* gene knockout Mab; Mab^ΔembC^::pVector, Mab^ΔembC^ with pMV261A’; Mab^ΔembC^::pembB^wt^, *embB^wt^* overexpressed in Mab^ΔembC^; Mab^ΔembC^::pembB^mutR350G^, *embB^R^*^350^*^G^*overexpressed in Mab^ΔembC^; Mab^ΔembC^::pembB^mutD306N^, *embB^D^*^306^*^N^*overexpressed in Mab^ΔembC^; Mab^ΔembC^::embB^edit-R350G^, *embB^R^*^350^*^G^*edited Mab^ΔembC^; Mab^ΔembC^::embB^edit-D306N^, *embB^D^*^306^*^N^*edited Mab^ΔembC^; Mab^ΔembC^::embB^mutR350G^, *embB^R^*^350^*^G^*spontaneous mutant Mab^ΔembC^; Mab^ΔembC^::embB^mutD306N^, *embB^D^*^306^*^N^*spontaneous mutant Mab^ΔembC^; Data are expressed as mean ± SD from three independent biological repeats. The experiment was performed in triplicate (three independent experiments), and the representative results are shown. **B)** Sensitivity of different Mab strains to crystal violet. The experiments were performed at least thrice, and only one representative image is shown in each case.

### Bioinformatics analysis of EmbB and EmbC

#### EmbB and EmbC proteins sequences comparison in Mab

Given that EmbB and EmbC express different arabinosyltransferases and are essential in cell wall synthesis, it was discovered in the experiment mentioned above that when the *embC* gene was absent, the *embB* gene mutation unexpectedly compensated the partially missing function of *embC*. Thus, we compared the amino acid sequences of EmbB and EmbC (Figure S1). EmbB and EmbC in Mab have 42.23% homology, and EmbB and EmbC confer the same amino acid at position 350, where EmbB showed the substitution from Arg to Gly in WGS results. The amino acid at the 555 position of EmbB is mutated to Ile as that in EmbC, and other mutations did not seem to be significantly related to the amino acid sequence of EmbC.

#### Amino acid sequences alignment of different mycobacterial proteins homologous to Mab-EmbB

Sequence comparison of the Mab^ΔembC^ spontaneous drug-resistant mutant gene found that the homologous proteins of Mtb, *M. bovis*, *Mycobacterium marinum* (*M. marinum*), *Mycobacterium avium* (*M. avium*), *Mycobacterium ulcerans* (*M. ulcerans*), and *Mycobacterium canettii* (*M. canettii*) are arabinosyltransferases B (EmbB). Sequence alignment showed that Mab-EmbB has about 67% identity with EmbB in Mtb, *M. bovis*, *M. avium*, *M. marinum*, *M. canettii*, and *M. ulcerans* (Table S1). The spontaneous resistance mutation sites are shown in Figure S2. It can be seen that the mutation sites are conserved sequences of mycobacterial EmbB, and mutations may change the function of EmbB.

## DISCUSSION

Mab, the most common NTM pathogenic bacteria, is naturally resistant to various antibiotics^3, 23^. Only a handful of antibiotics, such as CLA, AZT, AMK, etc., are effective in treating diseases caused by some but not all Mab^24^. Therefore, the discovery and development of new drugs are absolutely essential. Even though Mab’s pathogenicity has been widely studied, however, further investigation is still required to determine the precise mechanism of Mab resistance. Understanding the functionality of Mab proteins is crucial for avoiding external persecution.

Most of bacteria have a protective outer shell (cell wall), and Mab’s low permeability cell wall plays a vital role in their intrinsic resistance^25^. Arabinosyltransferase C, a protein involved in cell wall synthesis, is encoded by the *embC* gene, and the impairment of cell wall synthesis caused by the loss of *embC* increases the permeability of the cell wall in Mab, making the bacteria more sensitive to antibiotics^19, 26^.

In this study, we found that the MIC of ECH against Mab was 2 µg/mL, but it changed to 0.0078-0.0156 µg/mL against Mab^ΔembC^. The ECH resistance-related *embB* gene was identified by WGS of the spontaneously resistant Mab^ΔembC^ mutant strains, and the following mutations (Asp306Ala, Asp306Asn, Arg350Gly, Val555Ile, Gly581Ser) were found in EmbB. The *embB*, an essential gene that encodes arabinosyltransferase B is involved in cell wall synthesis. Overexpression of the *embB* gene in Mab^ΔembC^ could increase resistance to ECH (MIC: 0.25-0.5 µg/mL, 32 to 128-fold increased).

Simultaneously, the *embB* in Mab^ΔembC^ was edited using the CRISPR/Cpf1-assisted homologous recombination system developed by us recently^27,28^. These edited strains also became more resistant to ECH (MIC: 0.25-0.5 µg/mL). In addition, the protein sequence comparison exhibited a 42.23% similarity between the amino acid sequences of EmbB and EmbC. The above results suggest that *embB* and *embC* are associated with ECH-resistance in Mab. Considering the involvement of both EmbB and EmbC in cell wall synthesis^26^, we hypothesized that EmbB and EmbC may change the sensitivity of Mab to ECH by modifying cell wall permeability. Our hypothesis was confirmed through monitoring the EtBr accumulation by the *embB* overexpressed Mab^ΔembC^ and *embB* edited Mab^ΔembC^ strains. While drug-resistant Mab^ΔembC^ isolates with *embB* mutations were also acquired when screened with different antibiotics, such as LZD (our unpublished data). These findings suggest that when Mab^ΔembC^ is exposed to antibiotic stress, EmbB may mutate to compensate for the loss of partial function of EmbC to ensure its survival.

In conclusion, this study discovered that a natural product, ECH, has potent inhibitory activity against both Mtb and Mab. However, elucidation of the mechanism of ECH to reveal its potential targets in mycobacteria still needs to be outlined in detail. We found two homologous genes in Mab related to ECH resistance, *embB* and *embC*, encoding arabinosyltransferases and participating in the transport process of arabinosyltransferase. When EmbC in Mab is missing and unable to perform its normal function, EmbB will mutate under external pressure to compensate for the lost function of EmbC to adapt environmental changes. Our findings propose that Mab can change the sensitivity to ECH through functional compensatory mechanisms.

## MATERIALS AND METHODS

### Drug formulation

ECH was provided by the South China Sea Institute of Oceanology, Chinese Academy of Sciences, and dissolved in dimethyl sulfoxide (DMSO). MXF, LEV, VAN, RIF, RFB, LZD, CLA, CEF, CLF, apramycin (APR), and anhydrotetracycline (ATC) were bought from MeilunBio and dissolved in DMSO, whereas AMK (Meilunbio) and kanamycin (KAN) (Meilunbio) were dissolved in distilled water. Zeocin-Bleomycin (ZEO) (InvivoGen).

### Strains and growth conditions

Mab, AlMab, and Mab^ΔembC^ were cultured in Middlebrook 7H9 broth medium containing 10% Oleic Albumin Dextrose Catalase (OADC) and 0.05% Tween 80 at 37LJ for 3-5 days until they reached the expected concentrations or on 7H10/7H11 solid medium containing 10% OADC 37LJ for 7 days. *E. coli* trans-T1 (TransGen Biotech) was cultured at 37LJ in LB broth/solid medium with appropriate antibiotics.

The antibiotic concentrations (µg/mL) used for Mab and Mab^ΔembC^ were: KAN 100, APR 230, and ZEO 30, while for *E. coli*: KAN 50, APR 50, and ZEO 30.

### Testing the sensitivity of ECH against Mab and Mab^ΔembC^

#### *in vitro In vitro* activity of ECH against AlMab

AlMab was inoculated into 5 mL of 7H9 broth medium containing 10% OADC and 0.05% Tween 80. The culture was incubated at 37LJ and 200 rpm until the OD_600_ reached 0.6-0.8. RLUs were measured by a luminescence detector (Promega GloMax2020™) at 12 h intervals. If the RLUs of 200 μL bacteria was higher than 2 × 10^7^, then dilute the bacterial culture with 7H9 without Tween 80 until RLUs/ 200 µL dilution reaches around 3000-5000 as the tested culture.

ECH was diluted with DMSO to make the concentrations (mg/mL) of 1.6, 0.8, 0.4, 0.2, 0.1, 0.05, and CLA: 0.8 mg/mL. 2 µL of different concentrations of ECH was mixed with 198 µL of diluted AlMab in a 1.5 mL EP tube individually to get the tested concentrations (µg/mL): 16, 8, 4, 2, 1, and 0.5. Three replicates for each concentration were measured. Simultaneously, positive (AlMab with CLA 8 µg/mL) and negative (bacterial culture with DMSO) control groups were included in this assay. MIC_lux_ of ECH against AlMab was defined as the lowest concentration inhibiting more than 90% RLUs compared to that of the solvent control.

#### *In vitro* activity of ECH against Mab^ΔembC^

The activity of ECH against Mab^ΔembC^ was detected by broth dilution method: Mab^ΔembC^ was inoculated into 5 mL 7H9 broth medium and cultured at 37LJ and 200 rpm until OD_600_ reaches 0.6-0.8. Then, the culture was diluted to an OD_600_ of 0.125, and the ECH was diluted to 2 μg/mL with 7H9 without Tween 80. Finally, the working concentrations (μg/mL) of ECH were 1, 0.5, 0.25, 0.125, 0.0625, 0.03125, 0.0156, 0.0078, 0.0039, and 0. The diluted Mab strains were added into 96-wall plates with 2-fold serial drug dilutions and incubated at 37LJ for 7 days. MIC_liquid_ was defined as the minimum drug concentration at which the bacterial growth was not visible by the naked eye.

#### Screening of spontaneous ECH-resistant mutants using Mab**^Δ^**^embC^

The Mab^ΔembC^ was cultured in 100 mL of 7H9 liquid medium (containing 10% OADC, 0.05% Tween 80 and 5% ethyl bromide as mutagen) in a 250 mL conical flask and incubated at 37LJ and 220 rpm to obtain the OD_600_ of 0.6-0.8. 500 µL of ten-fold concentrated bacterial solution was plated on 7H10 plates containing different concentrations (μg/mL, 0.25, 0.5, 1, 2) of ECH and incubated at 37LJ. Three plates were set for each concentration. Each individual colony was picked and inoculated into 2 mL of 7H9 in a 50 mL tube and incubated at 37LJ and 200 rpm. The liquid culture was diluted and spread to obtain pure single colonies. Then the sensitivities of these isolated and pured colonies to ECH were detected.

#### Identification of the mutant gene and mutation sites

Mab^ΔembC^ spontaneous resistant mutants were randomly selected and sent to Genewiz Biology Company (Suzhou) after DNA extraction for WGS. The sequencing reads of Mab^ΔembC^ were compared with that of the parent strain to seek meaningful mutation sites. Primers were designed to amplify and validate the corresponding mutant genes of these strains.

#### Drug susceptibility testing of Mab**^Δ^**^embC^ mutants

The broth dilution method was used to detect the sensitivity of different antibiotics to Mab^ΔembC^ mutants. Stock solutions of ECH, RIF, RFB, MXF, LEV, LZD, CEF, VAN, and AMK were diluted in 7H9 without Tween 80 containing less than 2% DMSO. The Mab strains were cultured in 7H9 until OD_600_ reached 0.6-1 and then diluted to OD_600_ 0.125 with 7H9 without Tween 80. Then, 100 μL diluted culture and 100 μL drug were added into each well. The MIC_liquid_ was read out after continuous incubation at 37LJ for 7 days.

#### Overexpression of *embB* gene in Mab**^Δ^**^embC^

_Six versions (*embB*wild type, *embB*Val555Ile, *embB*Arg350Gly, *embB*Gly581Ser,_ *embB*^Asp306Ala^, *embB*^Asp306Asn^) of *embB* gene were amplified from Mab^ΔembC^ and spontaneous-resistant Mab^ΔembC^ mutants by PCR using primers CZ-Mab_embB-F/CZ-Mab_embB-R (5’-GGCCAAGACAATTGCGGATCCATGACAGAGAATTCCGTGACAGATAC-3’/5-ACATCGATAAGCTTCGAATTCTTACGGCTTGATCCGGATCTG-3’) and inserted into the pMV-261A’ plasmid under the control of the *hsp60* promoter. The six plasmids were transformed into the Mab^ΔembC^. The MICs of the recombinant strains to ECH were determined as described above.

#### *embB* gene editing in Mab^ΔembC^

Editing *embB* in Mab^ΔembC^ was performed through the CRISPR-Cpf1 assisted homologous recombination system as previously described^28^. Firstly, the pJV53-Cpf1 plasmid was electroporated into the Mab^ΔembC^ for Mab^ΔembC^::pJV53-Cpf1, which was then prepared as competent cells. The primers (Table 1) were designed to synthesize two single chains, which were annealed and linked with linearized pCR-Zeo to obtain pCR-embB^mut^. The 59 bp single-stranded DNA containing the mutated nucleotide(s) were synthesized as templates for creating point mutations. The five pCR-embB^mut^ plasmids and corresponding single-stranded template DNA were transformed together into the competent cells of Mab^ΔembC^::pJV53-Cpf1 via electroporation. The transformants were plated on 7H11 containing ATC 200 ng/mL, KAN 100 µg/mL, ZEO 30 µg/mL and incubated for 7 days. After confirmation of successful *embB* gene editing, the MICs_liquid_ of the edited strains to ECH were determined as described above.

**Table 8.**
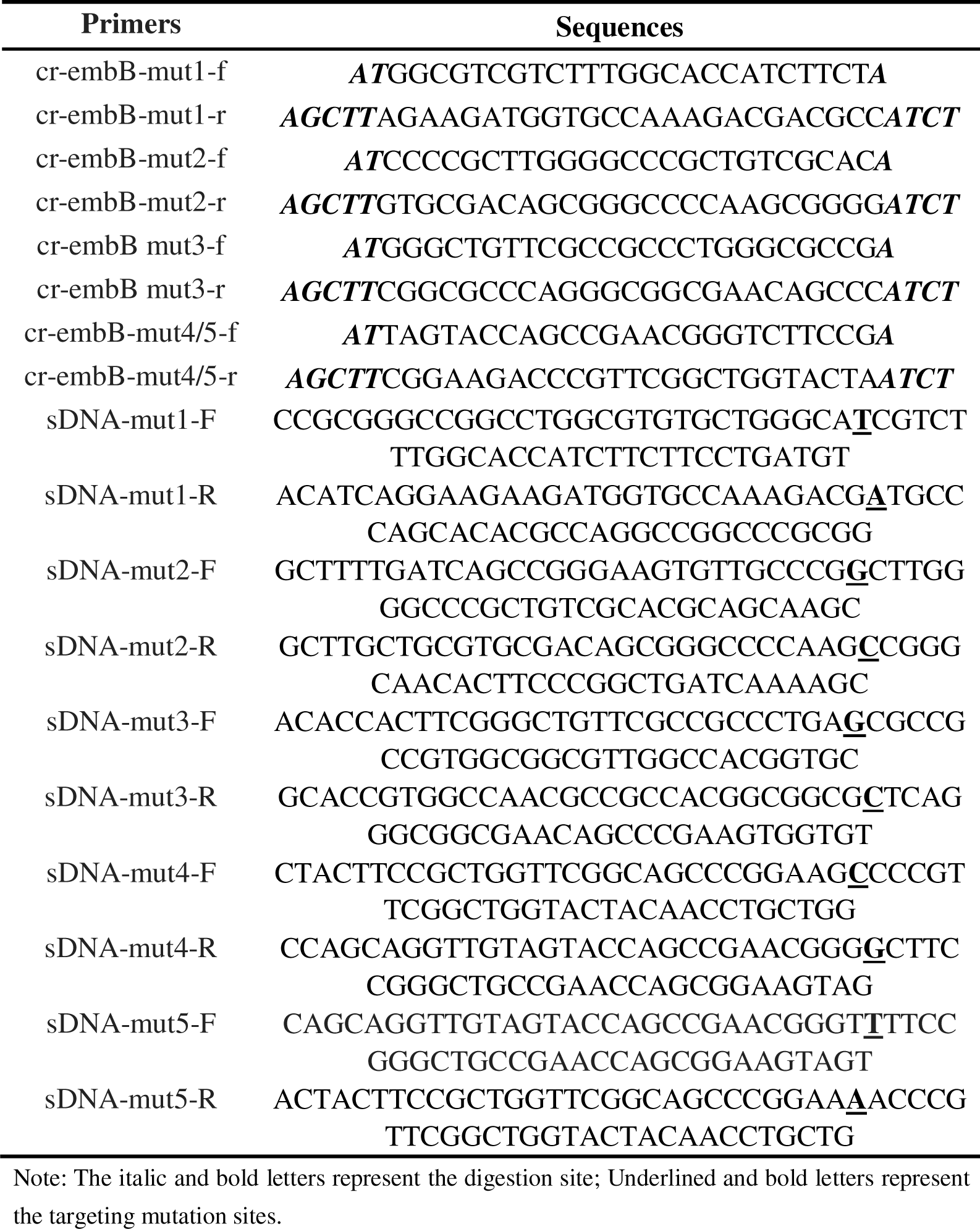
Primers used in gene editing.

#### Sensitivity of different strains to ECH

Detection sensitivities of the overexpressed and edited strains to ECH: The overexpressed and edited strains as well as Mab^ΔembC^ were cultured to reach the OD_600_ 0.6. 2 µL of the serial tenfold diluted culture (10^0^, 10^-1^, 10^-2^, 10^-3^, 10^-4^, and 10^-5^) was dropped in each square on 7H10 plates containing different concentrations of ECH and incubated at 37LJ for 7 days. At the same time, diluted the bacterial cultures to OD_600_ 0.05, and treated with different concentrations of ECH in triplicate. At the indicated time points, tested the CFU (colony forming unit) counts on 7H10 plates.

Detecting the sensitivity of different strains of the same mutant type to ECH: We selected a mutant type, *embB*^R350G^ and cultured Mab^ΔembC^, Mab^ΔembC^::embB^mutR350G^, Mab^ΔembC^::embB^editR350G^, Mab^ΔembC^::pVector, Mab^ΔembC^::pembB^mutR350G^, Mab^ΔembC^::pembB^wt^, in 7H9 broth at 37LJ and 200 rpm until OD_600_ 0.6-0.8. 2 µL of the serial tenfold diluted culture (10^0^, 10^-1^, 10^-2^, 10^-3^, 10^-4^, and 10^-5^) was dropped in each square on 7H10 plates containing different concentrations of ECH and incubated at 37LJ for 7 days. And as previously described, tested the CFU counts on 7H10 plates at the indicated time points.

#### Cell wall permeability detection

Two mutant types (*embB*^D306N^ and *embB*^R350G^) were selected to detect the effects on the cell wall permeability of Mab^ΔembC^ by EtBr accumulation testing^29^. Mab^wt^, _MabΔembC, MabΔembC::embBmutR350G, MabΔembC::embBmutD306N, MabΔembC::embBeditR350G,_ Mab^ΔembC^::embB^editD306N^, Mab^ΔembC^::pVector, Mab^ΔembC^::pembB^mutR350G^, Mab^ΔembC^::pembB^mutD306N^, Mab^ΔembC^::pembB^wt^, Mab^ΔembC^::pembC were grown until the OD_600_ reached 0.8. Bacteria were collected by centrifugation and washed with PBS-Tw (PBS with 0.05% Tween 80), and the OD_600_ was adjusted to 0.4. 1 mg/mL EtBr dissolved in DMSO was diluted into 4 μg/mL solution of PBS-Tw and the 0.8% glucose. 100 μL of diluted culture was added to each well of the white 96-well plate, and three replicates were set for each strain. A mixture of 100 μL diluted culture containing 4 μg/mL EtBr and 0.8% glucose was added into each well. Two control groups, sterile control (100 μL PBS-Tw + 100 μL EtBr) and non-EtBr control (100 μL PBS-Tw + 100 μL diluted bacteria) were arranged with three replicates per group. Real-time fluorescence of the 96-well plate was measured by a multifunctional microplate reader (PerkinElmer) with excitation and emission wavelengths set at 530 and 590 nm, respectively. The fluorescence values of the experimental group were subtracted from the EtBr baseline fluorescence values obtained in the corresponding sterile control group while ensuring that the fluorescence measurements of the non-EtBr control group were very low and did not increase over time.

#### Susceptibility to chemical compounds

We tested the toxicity of crystal violet (8 µg/mL) to different strains. Mab^wt^, Mab^ΔembC^, Mab^ΔembC^::embB^mutD306N^, Mab^ΔembC^::embB^editD306N^, Mab^ΔembC^::pVector, Mab^ΔembC^::pembB^mutD306N^, Mab^ΔembC^::pembB^wt^, Mab^ΔembC^::pembC were grown until the OD_600_ reached 0.8. 2 µL of the serial tenfold diluted culture (10^0^, 10^-1^, 10^-2^, 10^-3^, 10^-4^, and 10^-5^) was dropped in each square on 7H10 plates containing different concentrations of crystal violet and incubated at 37LJ for 7 days.

#### Bioinformatics analysis of EmbB and EmbC

Amino acid sequences of Mab-EmbB and Mab-EmbC from NCBI (https://www.ncbi.nlm.nih.gov/) were downloaded and compared with each other. Moreover, MAB_0185 (MAB-EmbB) homologous proteins were searched for in other mycobacteria, including Mab, Mtb, *M. bovis*, *M. avium*, *M. marinum*, *M. canettii*, and *M. ulcerans*, Sequence alignment was performed to analyze whether the mutants of Mab-EmbB were located in a conserved sequence.

## ACKNOWLEDGMENTS

This work was supported by the National Key R&D Program of China (2021YFA1300900, 2019YFA0901903), the National Natural Science Foundation of China (21920102003, 82022067, 22037006), the Chinese Academy of Sciences Grants (154144KYSB20190005, YJKYYQ20210026), the Key R&D Program of Sichuan Provenience (2023YFSY0047), the State Key Laboratory of Respiratory Disease, Guangzhou Institute of Respiratory Diseases, First Affiliated Hospital of Guangzhou Medical University (SKLRD-Z-202414, SKLRD-OP-202324, SKLRD-Z-202301, SKLRD-OP-202113 and SKLRD-Z-202412), Guangzhou Science and Technology Plan-Youth Doctoral “Sail” Project (2024A04J4273), President’s International Fellowship Initiative-CAS (2023VBC0015) and National Foreign Young Talent Program (QN2022031002L). We also acknowledge the group of Yicheng Sun from the Institute of Pathogenic Biology, Chinese Academy of Medical Sciences for kindly sending the pJV53-Cpf1 and pCR-Zeo plasmids as the tool of gene editing.

## AUTHOR CONTRIBUTIONS

**Jing He:** Investigation (lead); methodology (equal); software(equal); validation (equal); visualization (lead); writing-original draft (lead). **Yamin Gao:** Data curation (equal); formal analysis (equal); methodology (equal); writing-original draft (supporting); writing-review and editing (equal); funding acquisition (equal). **Jingyun Wang:** Investigation (supporting); data curation (equal). **H.M. Adnan Hameed:** Data curation (equal); writing-original draft (equal); writing-review and editing (lead); funding acquisition (equal); project administration (equal); supervision (equal). **Shuai Wang:** Conceptualization (supporting); formal analysis (equal); funding acquisition (supporting). **Cuiting Fang:** Methodology (supporting); data curation (equal). **Xirong Tian:** Investigation (supporting); validation (supporting). **Jingran Zhang:** Investigation (supporting); methodology (supporting) **Xingli Han:** Investigation (supporting); Software (supporting). **Yanan Ju:** Investigation (supporting); Software (supporting). **Yaoju Tan:** Supervision (supporting); funding acquisition (equal). **Junying Ma:** Resources (equal); funding acquisition (equal); project administration(equal). **Jianhua Ju:** Resources (equal); data curation (equal); funding acquisition (supporting). **Jinxing Hu:** Supervision (supporting); validation (equal). **Jianxiong Liu:** Conceptualization (equal); funding acquisition (equal); project administration (equal). **Tianyu Zhang:** Conceptualization (lead); funding acquisition (lead); project administration (lead); resources (supporting); validation (equal); writing-review and editing (equal).

## ETHICS STATEMENT

This study did not conduct animal or human experiments. There are no ethical issues involved.

## CONFLICTS OF INTEREST

The authors declare no conflict of interests.

## DATA AVAILABILITY STATEMENT

All data generated or analyzed during this study are included in this article.

## Notes

### Competing Interest Statement

The authors have declared no competing interest.

