## Supplementary Information for "EmbB and EmbC Regulate the Sensitivity of *Mycobacterium abscessus* to Echinomycin"

A

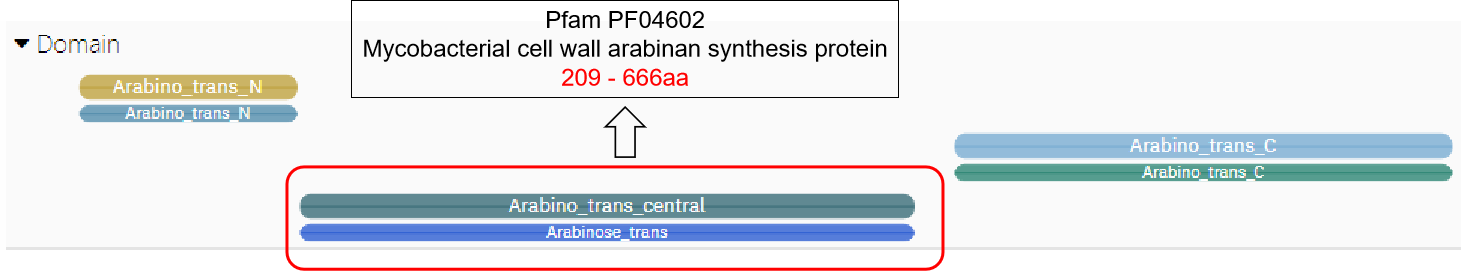

B

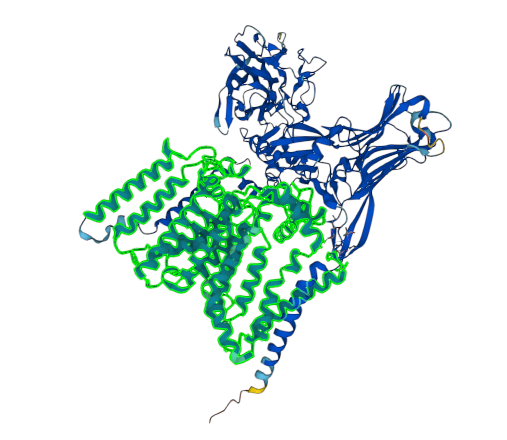

**Figure S1.** A) InterPro predicts the function domain of EmbB. B) The protein structure forecasted by Alphafold. The green displays the central structural domain.

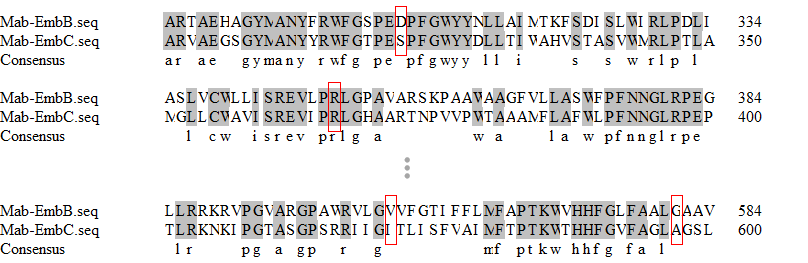

**Figure S2.** Partial results of Mab’s EmbB and EmbC proteins sequences alignment. The red boxes highlight the mutation sites.

**Table S1**. Homologous proteins of different mycobacteria and Mab-EmbB and their protein identities

| Strain | homologous protein | identity |
| --- | --- | --- |
| Mtb | arabinosyltransferase EmbB | 66.91% |
| *M. bovis* | arabinosyltransferase | 67.60% |
| *M. canettii* | arabinosyltransferase EmbB | 68.24% |
| *M. marinum* | putative arabinosyltransferase B | 67.23% |
| *M. avium* | arabinosyltransferase EmbB | 67.79% |
| *M. ulcerans* | arabinosyltransferase EmbB | 67.15% |

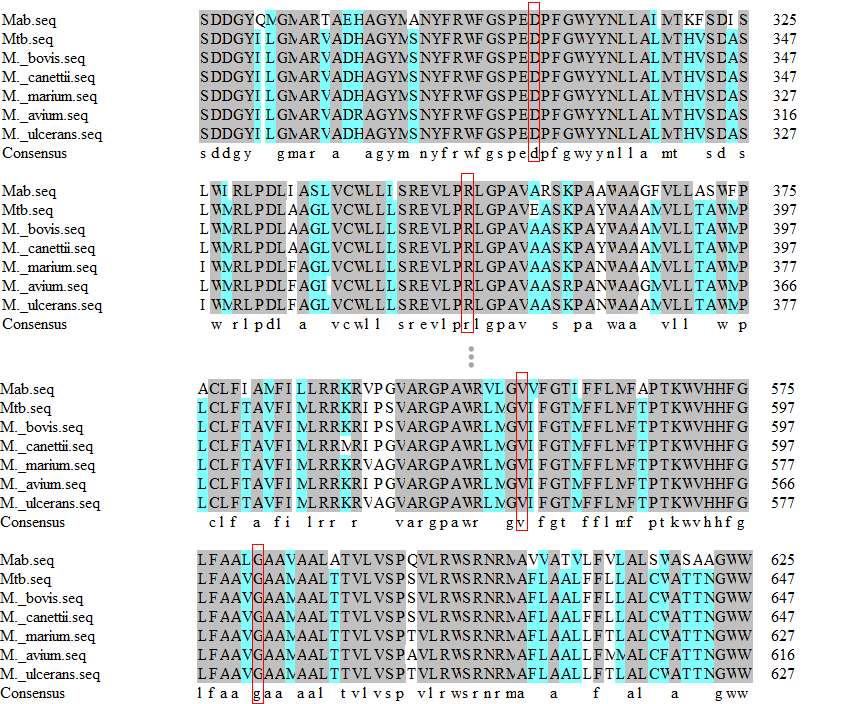

**Figure S2.** Partial sequence alignment of EmbB homologs in different mycobacteria and EmbB mutation positions of Mab. The red boxes highlight the mutation sites.
